# Focused ultrasound stimulates ER localized mechanosensitive PANNEXIN-1 to mediate intracellular calcium release in invasive cancer cells

**DOI:** 10.1101/2020.04.03.024372

**Authors:** Nan Sook Lee, Chi Woo Yoon, Qing Wang, Sunho Moon, Kweon Mo Koo, Hayong Jung, Ruimin Chen, Laiming Jiang, Gengxi Lu, Antony Fernandez, Robert H. Chow, Andrew C. Weitz, Paul M. Salvaterra, Fabien Pinaud, K. Kirk Shung

## Abstract

Focused ultrasound (FUS) is a rapidly developing stimulus technology with the potential to uncover novel mechanosensory dependent cellular processes. Since it is noninvasive, it holds great promise for future therapeutic applications in patients used either alone or as a complement to boost existing treatments. For example, FUS stimulation causes invasive but not noninvasive cancer cell lines to exhibit marked activation of calcium signaling pathways. Here, we identify the membrane channel PANNEXIN1 (PANX1) as a mediator for activation of calcium signaling in invasive cancer cells. Knockdown of PANX1 decreases calcium signaling in invasive cells, while PANX1 overexpression enhances calcium elevations in non-invasive cancer cells. We demonstrate that FUS may directly stimulate mechanosensory PANX1 localized in endoplasmic reticulum to evoke calcium release from internal stores. This process does not depend on mechanosensory stimulus transduction through an intact cytoskeleton and does not depend on plasma membrane localized PANX1. Plasma membrane localized PANX1 however plays a different role in mediating the spread of intercellular calcium waves via ATP release. Additionally, we show that FUS stimulation evokes cytokine/chemokine release from invasive cancer cells, suggesting that FUS could be an important new adjuvant treatment to improve cancer immunotherapy.

## INTRODUCTION

Cancer cell invasion and tumor metastasis play a critical role in cancer mortality. The mechanisms by which malignant tumors leave the primary tumor site, invade, and metastasize to other organs are complex, interrelated and only partially understood. Calcium signaling, however is known to be critical in these processes. To develop a functional assay of cancer cell invasion potential, we recently used FUS stimulation to probe the altered calcium signaling pathways exhibited by invasive cancer cells (Hwang et al., 2013; Weitz et al., 2017). FUS stimulation caused invasive, but not noninvasive cancer cell lines, to exhibit marked calcium signaling suggesting a novel means to determine the invasion potential. We validated this using a Matrigel invasion assay, demonstrating that the degree of invasion correlates well with the degree of FUS-dependent Ca^2+^ signaling (Hwang et al., 2013; Weitz et al., 2017). FUS stimuli evoke widespread Ca^2+^ oscillatory dynamics in several invasive cancer cell lines (breast MDA-MB-231, prostate PC-3 and bladder T24/83), but not in noninvasive cells of the same cancer type (MCF-7, BPH-1, and RT112/84) suggesting that this is a general property of invasive cells (Hwang et al., 2013; Weitz et al., 2017). Also, different FUS stimulation frequencies result in similar responses indicating that Ca^2+^ signaling is independent of stimulation frequency (3-, 38- or 200-MHz).

FUS stimulation of invasive cells also results in a time dependent propagation of an extracellular calcium wave spreading away from cells located at the transducer focus. The mechanism(s) for extracellular calcium wave propagation is unclear, it does not depend on ultrasonic surface waves or gap junctions (Weitz et al., 2017). Additional pharmacological studies in invasive cancer cells suggested the involvement of <u>IP_3_</u> receptors (IP_3_Rs) or TRP channels (Weitz et al., 2017), as suggested by others (Bootman et al., 2002; Diver et al., 2001; Xu et al., 2005). Non-focused US also stimulates calcium signaling mediated by the Piezo1 mechanosensitive ion channel directly couple to microbubbles (Pan et al., 2018). Other ER-localized mechanosensitive channels (i.e. Msy1 and Msy in fission yeast) regulate intracellular Ca^2+^ and cell volume for survival upon hypo-osmotic shock (Nakayama et al., 2012).

FUS technology has also been proposed for use in cancer therapy and particularly immunotherapy. High-intensity (>5 W/cm^2^) continuous FUS generates a systemic immune stimulatory effect resulting in tumor ablation (Lu et al., 2009). Pulsed FUS (i.e., non-continuous stimulus to minimize heat generation) (Hersh et al., 2016) may induce a more refined cellular/molecular immune response, (Ziadloo et al., 2012) by initiating inflammatory responses which boost cancer immunotherapy (Curley et al., 2017; Mauri et al., 2018). FUS may thus offer a new approach to overcome cancer immune-resistance, a well-known limitation preventing more wide-spread clinical adoption of successful immunotherapies such as CAR T cells (Caliendo et al., 2019; Tokarew et al., 2019). A detailed understanding of the mechanistic aspects of FUS-response mechanisms however is currently lacking.

In this study we establish a new role for the mechanosensitive PANX1 hemichannel (Bao et al., 2004) in mediating Ca^2+^ signaling in invasive cancer cells. PANX1 localizes to both plasma membrane (PM) as well as endoplasmic reticulum (ER) (Vanden Abeele et al., 2006). Mechano-sensitive responses have previously been described in neurons, including retinal ganglion cells (Xia et al., 2012) and other cell types. Our results suggest that FUS can directly stimulate ER localized PANX1 in invasive PC-3 cancer cells to generate Ca^2+^ release from intracellular ER stores, independently of extracellular Ca^2+^ entry. This is a newly described role of PANNEXIN-1 as a regulator of calcium ion exchange between the ER and cytoplasm, suggesting a new working model of how FUS interacts with cancer cells to initiate and propagate Ca^2+^ signaling. In addition, our results suggest that continued development of FUS technology could provide not only a new way to probe mechanosensitive functions of signaling pathways located in specific intracellular compartments but also to harness the potential to regulate adjunct immune cell responses through the coupling of mechanosensory stimulus to chemokine/cytokine release profiles.

## MATERIALS AND METHODS

### Cell Lines

PC-3 prostate cancer lines and HEK 293T cells were used in this study. PC-3 cells were purchased from ATCC and HEK 293T cells obtained from Dr. Fabien Pinaud at University of Southern California. Both cells were cultured in DMEM. The medium was supplemented with 10% FBS and 2 mM l-glutamine, and P/S. All cell lines were tested to be free of mycoplasma contamination using a mycoplasma PCR detection kit (Sigma). Cell lines were authenticated using short tandem repeat (STR) analysis by the University of Arizona Genetics Core. PC-3 cells are highly invasive cell lines, while HEK cells were used as non-invasive because of poor transfection efficiency of BPH-1.

### Cell Preparation and Transfection

Cells were plated on 35-mm culture dishes, or 24-well culture plates to a density of 10^6^ or 10^5^ cells per dish or well. All cells were stained with cell membrane permeant Fluo-4 AM (Thermo Fisher Scientific), a fluorescent reporter of intracellular calcium activity. Staining was performed by incubating dishes/wells with 1 µM Fluo-4 AM for 30 min immediately prior to imaging. Following calcium dye loading, cells were washed with and maintained in external buffer solution consisting of 140 mM NaCl, 2.8 mM KCl, 1 mM MgCl2, 2 mM CaCl2, 10 mM HEPES, and 10 mM d-glucose, adjusted to pH 7.3 and 290-300 mOsm.

Lipofectamine 3000 (Invitrogen) was used for cDNA construct transfection experiments. cDNA constructs used are WT PANX1-EGFP and mt PANX1-mRFP that are provided by Dr. Tavazoie (Furlow et al., 2015)]. FL WT PANX1 with no EGFP fusion (WT PANX1) was made for Fluo-4 AM Ca^2+^ detection, after deletion of EGFP using a standard sub-cloning method. For siRNA transfection, we used DharmaFect (Thermo Scientific) according to the manufacturer’s instructions. siRNA specifically designed to target FL PANX1 (si-*PANX1*) was purchased from Dharmacon (GE Healthcare) (D-018253-02) (Figure S 5). Control siRNA (negative) was obtained from GE Healthcare (D-001810-10).

### Analysis of mRNA expression

Expression of mRNA was quantified by qPCR as described previously(Lee et al., 2015). Primers are the followings: for *PANX1*, forward 5’-agcccacggagcccaagttca and reverse 5’-gcgcgaaggccagcgaga, for *GAPDH* and *CyclophilinA*, they are described in our previous paper(Lee et al., 2015).

### Ultrasound Transducers

A single-element, lithium niobite (LiNbO3), press-focused 46 MHz (f-number = 2, focal length = 6 mm) transducer was fabricated in house as described previously(Lam et al., 2013) and used in most experiments. In addition, a PZT, pressed-focused 3-MHz transducer (f-number = 1.5, focal length = 4 mm) was also tested. To drive the transducers, sinusoidal bursts from a signal generator (SG382; Stanford Research Systems) were fed to a 50-dB power amplifier (525LA; Electronics & Innovation) whose output was used to excite the transducer. For the 46-MHz transducer, amplitude was tested at different input Vp–p, pulse repetition frequency (PRF) at 1 kHz, and duty cycle at 5%, in PC-3 and HEK cells (Supplementary Figures 1 & 3). The acoustic output of the 46-MHz transducer was measured with a needle hydrophone (HGL-0085; Onda). Figure 1A shows the measured beam width of the focused ultrasound to be ∼70 μm, which may focus on ∼6 cells. Using the standard cell stimulation parameters provided above [12 Vp-p (40 mV) amplitude, 1 kHz PRF, and 5% duty cycle), the intensity and pressure at the focus were measured by the hydrophone.

**FIGURE 1.**
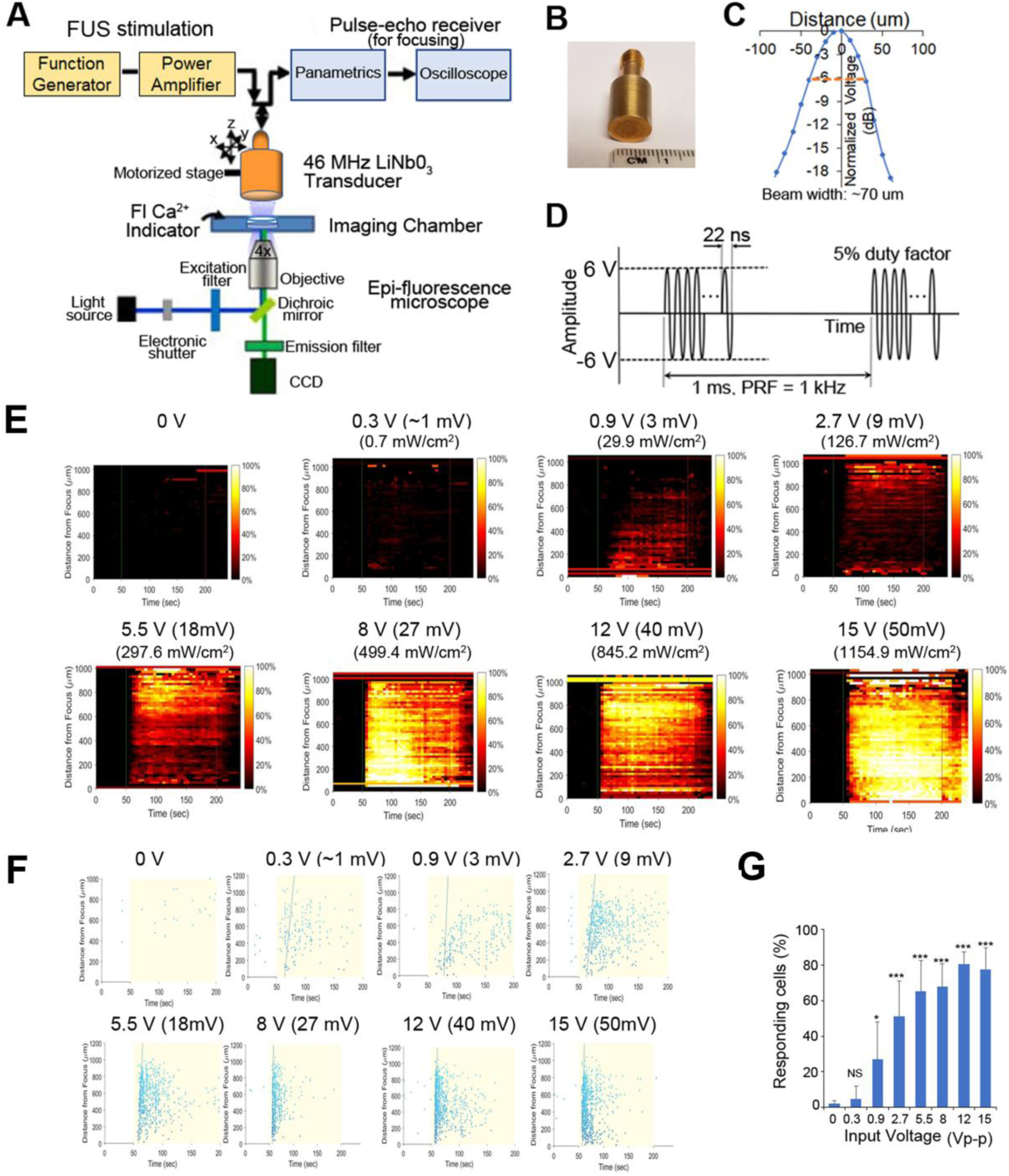
Schematic of our experimental system and effect of FUS stimulation amplitude on PC-3 Ca^2+^ response. **(A)**. A 46-MHz, single-element, LiNbO_3_, press-focused transducer was used for FUS stimulation and focused with a pulse-echo receiver. Cells were imaged using epi-florescence microscopy in the presence of a Ca^2+^ indicator. **(B)** Photograph of the 46-MHz transducer used in most experiments. **(C)** The beam width produced by the transducer was measured by hydrophone and was ∼70 μm. **(D)** Typical voltage waveform used to drive the transducer. Carrier frequency had an amplitude of 46 MHz (12 Vp–p), pulse repetition frequency was 1 kHz and duty cycle was 5%. **(E-F)** Effect of FUS stimulation amplitude on PC-3 Ca^2+^ response. Standard stimulus parameters (**D**) were used while varying the transducer input voltage. All stated voltages represent peak-to-peak amplitude (Vp-p). Values in parentheses indicate the mV and Ispta at each voltage, as measured by a hydrophone. **(E)** 2-D histograms showing the percentage of responding cells over time. **(F)** Scatter plots showing the time at which each individual cell first responds. **(G)** Quantitative percentage of responding cells. *n*>3 biological replicates. Error bars, SEM., *, P<0.05; **, P<0.01; ***, P<0.001 by a one-tailed t-test. *n* represents biological replicates.

### Ultrasound Stimulation and Fluorescence Imaging

A custom microscope system was used to image cellular fluorescence while performing simultaneous ultrasonic stimulation as described previously(Hwang et al., 2013; Weitz et al., 2017). Petri dishes or plates containing cells were placed on an inverted epifluorescence microscope (Olympus IX70), and the ultrasound transducer was lowered into the external buffer solution. A motorized three-axis micromanipulator was used to position the transducer in focus with the cell monolayer. In each experiment, live-cell fluorescence imaging was performed for 240 s (and sometimes, 300 s), with the ultrasound stimulus being delivered continuously between t = 50 and 200 s. Excitation light was provided by a mercury arc lamp and filtered through an excitation bandpass filter (488 ± 20 nm). Fluorescence emitted from the calcium dye was filtered through an emission bandpass filter (530 ± 20 nm) and recorded at 1 Hz (30% exposure duty cycle) with a digital CMOS camera (ORCA-Flash2.8; Hamamatsu). All imaging was performed at 4× magnification in order to capture activity from hundreds or thousands of cells simultaneously. For each cell line, simulation and imaging experiments were replicated in at least two different dishes of cells, and over least three independent fields of view per dish. Experiments involving pharmacological blockers were limited to a single field of view per dish. Figures show representative data obtained from one field of view.

### Data Processing

Data were post-processed to determine the calcium response of every imaged cell as described previously(Weitz et al., 2017). Cell locations were identified automatically with CellProfiler image analysis software(Carpenter et al., 2006) and used to extract the raw fluorescence intensities of each cell. These intensities were exported to MATLAB (MathWorks) in order to calculate each cell’s normalized change in fluorescence (ΔF/F) during every imaging frame. Responding cells were defined as those that exhibited a ΔF/Fmax greater than 3.5 times the pre-stimulus root-mean-square noise level. Two types of plots were generated for each 240 s experiment: a histogram showing the percentage of responding cells over time and a scatter plot indicating the time at which each cell first responded to the stimulus. Responding cells in these plots were arranged with respect to their distance from the transducer focus. The cell response index (CRI) was obtained as described previously(Hwang et al., 2013).

### Pharmacology

To investigate the mechanism of ultrasound-induced calcium rise in invasive cancer cells, PC-3 cells were stimulated in the presence of various pharmacological agents. We tested several different blockers, each applied separately (Table S1). Blockers were dissolved in the external buffer solution 15–30 min before performing imaging and ultrasound stimulation. Cellular responses were measured before adding the blockers and in the presence of blockers.

### ATP release assay

Cells were seeded in quadruplicate at 100,000–200,000 cells per well in 24-well plates and grown overnight. Each well was then washed with 1 ml external buffer solution (EBS). For PANX1 inhibition, cells were incubated at room temperature for 10 min in EBS supplemented with one of the following reagents: CBX (500 µM), probenecid (2 mM; Life Technologies), ^10^Panx1 (100 µM) or an equivalent dose of the appropriate vehicle control (EBS or scrambled peptide). The wash or pretreatment solution was then aspirated, replaced with 1 ml EBS for 10 min, collected and transferred to microcentrifuge tubes, and then spun at 86g for 2 min at room temperature. 50 or 100 μl of supernatants was transferred to 96-well plates and ATP was measured using the CellTiter-Glo Luminescent Cell Viability Assay (Promega) according to the manufacturer’s instructions.

### Immunofluorescence and confocal microscopy

Cells expressing fluorescently tagged proteins were fixed in 4% paraformaldehyde, stained with DAPI (Thermo Fisher) and mounted using ProLong Gold antifade reagent (Thermo Fisher), and imaged using Leica TCS microscope.

### TIRF imaging and Immunocyto staining (ICS)

TIRF microscopy images were acquired on an inverted Nikon Eclipse Ti-E microscope, equipped with a 100× 1.49 NA objective (Nikon), an iXon EMCCD camera (Andor), laser lines at 405, 488, 561, and 647 nm (Agilent), a multiband pass ZET405/488/561/647× excitation filter (Chroma), a quad-band ZT405/488/561/647 dichroic mirror (Chroma), and appropriate emission filters for imaging of mRFP (600/50 nm, Chroma) and GFP (525/50 nm, Chroma). Illumination was performed by TIRF to ensure exclusive illumination of the plasma membrane.

For ICS, the cells are fixed with 100% methanol at -20°C for 10 min. After washing with PBS, they were permeabilized with 1% Triton X-100 at 37°C for 30 min. After blocking, add primary C-terminus anti-PANX1 (Santa Cruz Biotech.) or N-terminus anti-PANX1 (Alomone Labs) was added and incubated at 4°C at 12 hours followed by addition of second anti-mouse-PE (Santa Cruz) at room temperature for 30 min. After washing, the cover slips containing cells were mounted and observed using a confocal microscope (Leica).

### Human Cytokine assay

1 day after FUS stimulation, cell culture supernates were collected and centrifuged. 700 μl of supernates are applied for human XL cytokine array (R&D systems) according to the manufacturer’s instructions. Pixel densities on developed X-ray film were collected and analyzed using a transmission-mode scanner and image analysis software (Image Studio Lite).

### Statistical analyses

In general results are expressed as mean±s.e.m. Statistical analysis of multiple groups used one-way ANOVA, and Dunnet’s correction for multiple comparisons (GraphPad Prism, V8). Two group comparisons were tested using the Student’s t-test (one- and two-tailed) in Excel (v2016).

## RESULTS

### FUS stimulation evokes Ca^2+^ signaling in invasive PC-3 cells

To further clarify the mechanism(s) of FUS-dependent Ca^2+^response, our usual stimulus protocol used a 46-MHz, single-element, LiNbO_3_, press-focused transducer focused via a pulse-echo receiver coupled with epi-fluorescence microscopy to assess both intra- and intercellular changes in Ca^2+^ dynamics (**Figure 1A-D**, see also Methods for additional details). In our previous work we demonstrated that the magnitude of the FUS-induced Ca^2+^ response did not depend on the frequency of stimulation (Weitz et al., 2017). Here we examine its dependence on stimulus amplitude (i.e. intensity). Increasing voltage during FUS stimulation of PC-3 cells results in a larger Ca^2+^ response (i.e. a dose–response relationship as shown in **Figure 1E-G**). Standard stimulus parameters (see Materials and Methods, **Figure 1**) were used while varying the transducer input voltage (**Figure 1E, F**). Stimulation at ∼1 Vp–p, 2.7 Vp-p, and 12 Vp-p evoked calcium activity in ∼20%, ∼50%, and ^>^80% of cells, respectively (**Figure 1G**). We stimulated at 12 Vp-p for the remainder of this study, while keeping pulse repetition frequency (PRF) at 1 kHz and duty cycle at 5%. A 3-MHz stimulus was also effective in PC-3 cells (**Figure S1**), reconfirming the independence of stimulus frequency in eliciting Ca^2+^ responses (Weitz et al., 2017).

Our previous studies (Weitz et al., 2017) suggested ER localized IP_3_ receptors or PM localized TRP channels involvement in mediating invasive cancer cell FUS-dependent Ca^2+^ responses.

Here we use PC-3 cells as a model of an invasive cancer cell type and compared FUS-dependent Ca^2+^ responses to those in a non-invasive HEK293 cell line. Non-responsive HEK cells were chosen as an appropriate control line in this study, rather than previously used BPH-1 cells, since they were much easier to transfect than BPH-1 (∼90% vs. <5% transfection efficiency). As expected, FUS stimulation evoked strong Ca^2+^ responses in PC-3 cells but not in non-invasive HEK cells (**Figure 2A, B, Figure S2, Videos S1-S2**). In PC-3 cells, three distinct stimulus-dependent Ca^2+^ patterns are observed in individual cells in the presence of normal external Ca^2+^: Ca^2+^ oscillation, double Ca^2+^ spikes or a single spike (**Figure 2C**). We tested if PC-3 responses were mediated by Ca^2+^ influx by severely reducing or eliminating extracellular Ca^2+^ (0 or 20 µM vs. normal 2 mM). FUS stimulation in low or no external Ca^2+^ still exhibited Ca^2+^ response, but only a single spike pattern, in PC-3 cells (**Figure 2D**, Bottom). We additionally investigated Ca^2+^ influx blockers to assess their effects on FUS-dependent Ca^2+^ dynamics. Surprisingly, treatment of PC-3 cells with two different Ca^2+^ influx blockers (BTP2 or SKF96365) still showed all three patterns of Ca^2+^ response (**Figure S3, Table 1**) rather than the single spike when external Ca^2+^ is absent or low. This suggests that the specific route of Ca^2+^ entry may determine the specificity of subsequent response patterns. Notably, FUS stimulation in normal external Ca^2+^ (i.e. Ca^2+^ influx) following thapsigargin (TG) treatment completely abolished all Ca^2+^ responses (**Figure 2E**). TG is an agent that depletes intracellular Ca^2+^ stores. Our results indicate that external Ca^2+^ influx is not necessary for a FUS induced single spike Ca^2+^ response in PC-3 cells, suggest that this response is likely due to release from an internal storage site and differs from pharmacologically blocking 2 different PM Ca^2+^ channels. The mechanism of Ca^2+^ entry may thus play an important role in mediating complex Ca^2+^ dynamics following mechanosensory stimulation.

**Table 1.** Pharmacological effects of agents on FUS-stimulated Ca^2+^ dynamics in PC-3 cells.

| Agents | Target Effect | $\text{Ca}^{2+}$ Response |
| --- | --- | --- |
| **External $\text{Ca}^{2+}$ influx** |  |  |
| BTP2 | CRAC inhibitor, Blocks $\text{Ca}^{2+}$Influx | Normal, Figure S3 |
| SKF96365 | TRP antagonist, Blocks $\text{Ca}^{2+}$Influx | Normal, Figure S3 |
| **Intracellular $\text{Ca}^{2+}$ release** |  |  |
| Carbenoxolone | PANX1 inhibitor | Blocked, Figure S4 |
| Flufenamic acid | PANX1, CX43 inhibitor | Blocked, Figure S4 |
| Probenecid | PANX1 inhibitor | Blocked, Figure S4 |
| $^{10}\text{Panx1}$ | PANX1 inhibitor | Partly Blocked, Figure 3 |
| Xestospongine C | $\text{IP}_3\text{Rs}$ inhibitor | Partly Blocked, Figure 3 |
| **Interacellular $\text{Ca}^{2+}$ wave propagation** |  |  |
| Apyrase | Extracellular ATPase | Blocked, Figure 7 |
| Suramin | P2 purinergic receptor antagonist | Blocked, Figure 7 |
| PPADS | P2 purinergic receptor antagonist | Blocked, Figure 7 |
| AZ11645373 | Selective P2X7 inhibitor | Normal, Figure S3 |
| MRS2179 | Selective P2Y1 antagonist | Normal, Figure S3 |

**FIGURE 2.**
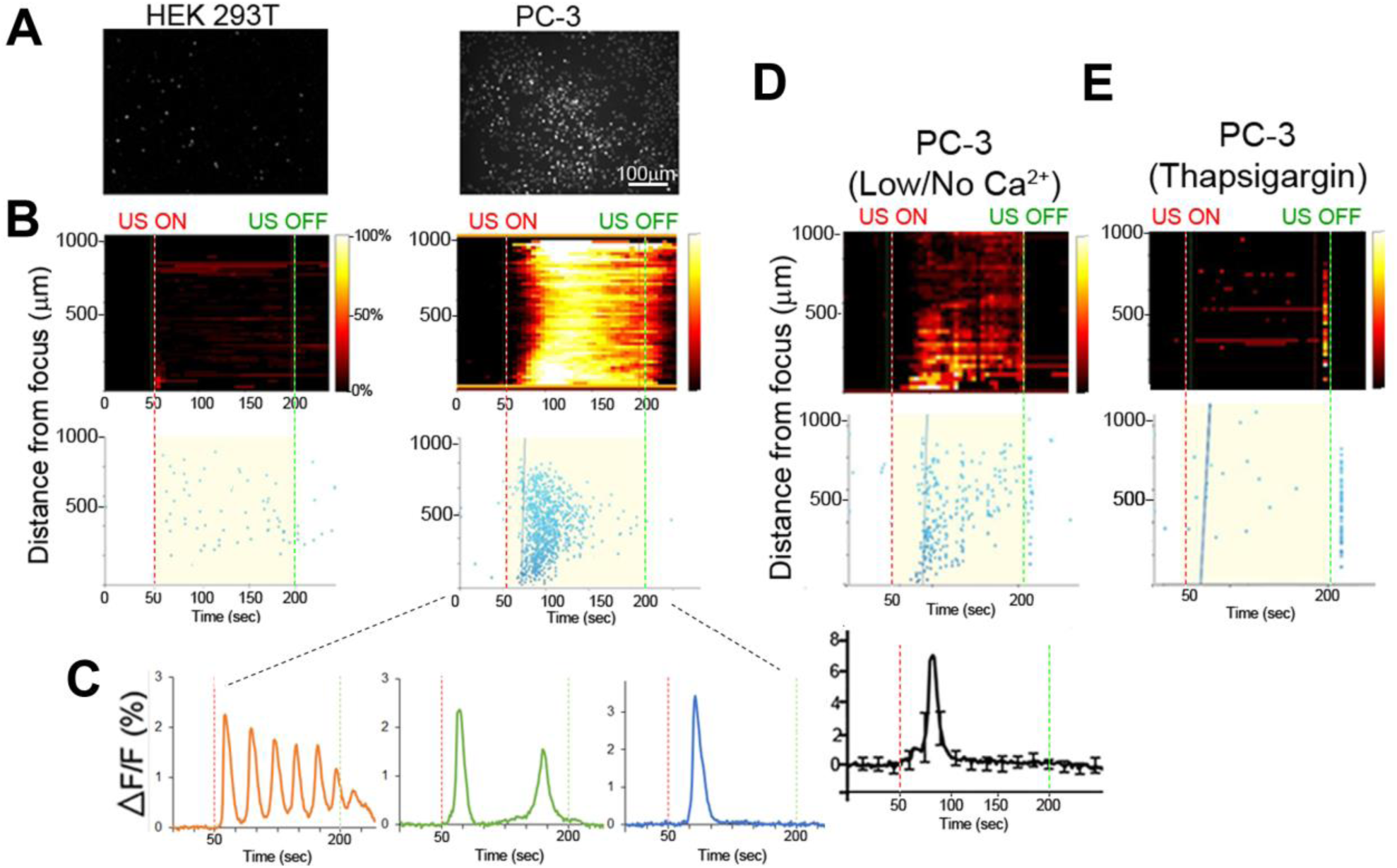
Ca^2+^ dynamics in invasive and non-invasive FUS stimulated cancer cells. **(A)** Background-subtracted fluorescence images show strong Ca^2+^ signaling in invasive PC-3 (left) but not non-invasive HEK (right) cells. **(B)** Top, 2-D histograms showing the percentage of responding cells over time. Vertical red and green dotted lines indicate FUS stimulus onset (50 s) and offset (200 s) times, respectively. Bottom, scatter plots showing the time of the first response in individual cells following stimulus. **(C)** Typical Ca^2+^ responses in invasive PC-3 cells exhibit either an oscillating (left), double (center) or single (right) spike pattern. **(D)** Ca^2+^ responses are present in PC-3 cells in external no or 20 mM (low) Ca^2+^ concentration. **(E)** Thapsigargin (TG) treatment in the normal external Ca^2+^ concentration (2 mM) drastically reduces the Ca^2+^ response.

### PANX1 mediates intracellular Ca^2+^ release in PC-3 cells

To further investigate the complex pattern of Ca^2+^ signaling following FUS stimulation, we tested Ca^2+^ release from an internal storage site. IP_3_ Receptors are known to be mediate Ca^2+^ release from ER or sarcoplasmic reticulum (SR) stores (Mery et al., 2005; Rizaner et al., 2016; Sasse et al., 2007). Treatment of cells with Xestospongin C (an IP_3_R inhibitor) partly inhibits the FUS-induced Ca^2+^ response (**Figure 3**). This suggests that other Ca^2+^ channels may also mediate release form internal stores. One potential candidate for such a role is PANX1. PM localized PANX1 is well studied for its role in ATP release but PANX1 is also localized to the ER where its function(s) is unknown (except for involvement in Ca^2+^ leaks, (Vanden Abeele et al., 2006)).

**FIGURE 3.**
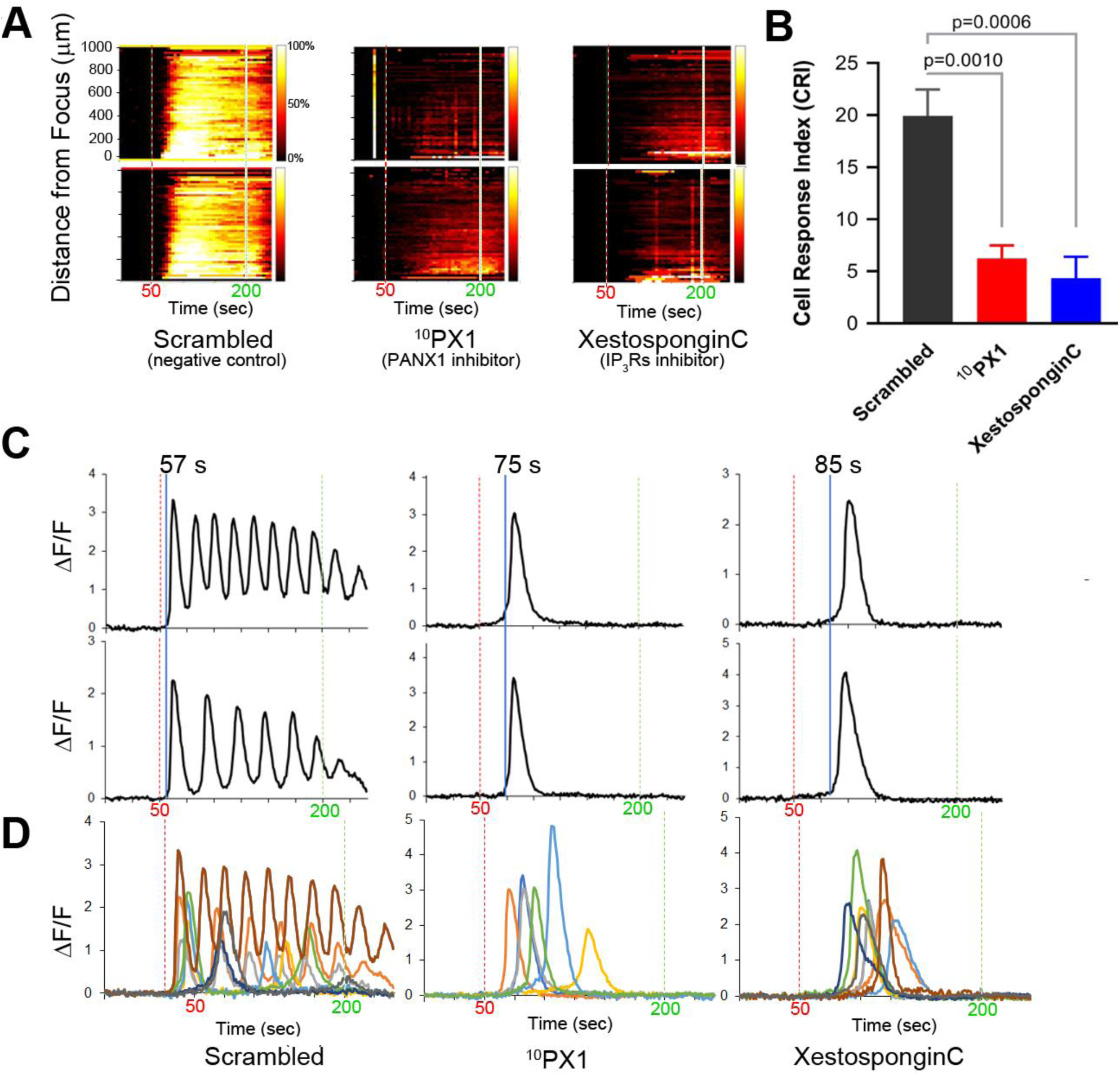
Treatment of PC-3 cells with ^10^PX1 (PANX1 inhibitor) abolishes the normal FUS-induced Ca^2+^ oscillation response but uncovers single Ca^2+^ transients. **(A)** Left column, the cells exhibited strong Ca^2+^ responses at 20 min after 200 μM scrambled peptide application as a control. Center column, cells were stimulated at 20 min after 200 μM ^10^Panx1 peptide (^10^PX1) application, and the responses were partly reduced. Right column, 20 min after 2 μM Xestospongin C (XC) application, the responses were also partly reduced. Two representative cells were shown in each treatment. **(B)** Quantitative CRI values of the inhibitor treatments. *n*=3 (XC), or *n*=6 (SC, ^10^PX1). Error bars, s.e.m., ANOVA, Dunnet’s correction, exact p values. **(C)** Fluorescence patterns in cells that first responded to the stimulus after the treatments. Two representative cells are shown by ΔF/F. **(D)** Fluorescence patterns in several cells that first responded to the stimulus after the treatments; Scrambled (9 cells), ^10^PX1 (5 cells) and XC (6 cells).

We treated of PC-3 cells with ^10^Panx1 peptide, a PANX1 inhibitor (Furlow et al., 2015). This results in a decrease in the FUS stimulated Ca^2+^ response (**Figure 3**). Ca^2+^ oscillations and double transients were eliminated but not the single transients (**Figure 3C, D**). This result is remarkably similar to Xestospongin C treatment (**Figure 3A-D**; see also **Videos S3-S5**). The primary difference between ^10^Panx1 and Xestospongin C-treated cells was in the timing of the single Ca^2+^ transients. ^10^Panx1treated cells had a ∼20 s delay after stimulation to onset (relative to the control) (**Figure 3C, D**, middle) while Xestospongin C had a slightly longer ∼30 s delay. In both cases the response was maintained for ∼30-40 s (**Figure 3C, D**, right). Treatment with a scrambled version of ^10^Panx1 exhibited the normal Ca^2+^ response (three patterns). Treatment with 2 additional PANX1 inhibitors probenecid and carbenoxolone (CBX) completely eliminated Ca^2+^ response (**Figure S4, Table 1**). These data indicate that both PANX1 and IP_3_Rs likely initiate and maintain FUS-induced Ca^2+^ oscillatory responses. Simultaneous addition of ^10^Panx1 and Xestospongin C did not further reduce the Ca^2+^response suggesting that the underlying mechanisms are complementary rather than independent (**Figure S4**), suggesting that they may be part of the same response pathway.

We next investigated a role for ER (or SR) localized PANX1 as a mediator of Ca^2+^ release by reducing PANX1 expression in PC-3 cells using si-*PANX1*RNA knockdown. Following treatment FUS-induced Ca^2+^ oscillations were variably reduced ∼50-70% (**Figure 4A, B**). Single or double calcium transients were delayed by ∼20s compared to the control response (**Figure 4C, Videos S6-S7**). We additionally transfected non-FUS responsive HEK cells (see **Figure 1E, F**) with WT PANX1 (i.e a full-length WT PANX1^1-425^ sequence). This construct had no EGFP fusion since this interferes with the Ca^2+^ imaging assay (see Methods). The WT PANX1 transfection converts HEK cells to robust FUS stimulation dependent Ca^2+^ responsiveness (**Figure 4D, E**). We also transfected HEK cells with a mutant form of PANX1^1–89^ lacking the normal C-terminal amino acids which was fused to mRFP (mt PANX1-mRFP). Interestingly, mt PANX1-mRFP transfection resulted in spontaneous Ca^2+^ activity even before FUS stimulation as well as exhibiting robust FUS-induced Ca^2+^ responsiveness (**Figure 4D, E**; **Videos S8-10**). The FUS-induced response in mt PANX1-mRFP transfected cells however was reduced relative to WT PANX1-transfected HEK cells (**Figure 4E**). Taken together these results indicate that PANX1 appears to be both necessary to generate FUS dependent Ca^2+^ responsiveness in PC-3 cells and sufficient to convert non-responsive HEK cells to a responsive state as well as generating non-FUS Ca^2+^ internal release dependent (i.e. Ca^2+^ leaks (Vanden Abeele et al., 2006)).

**FIGURE 4.**
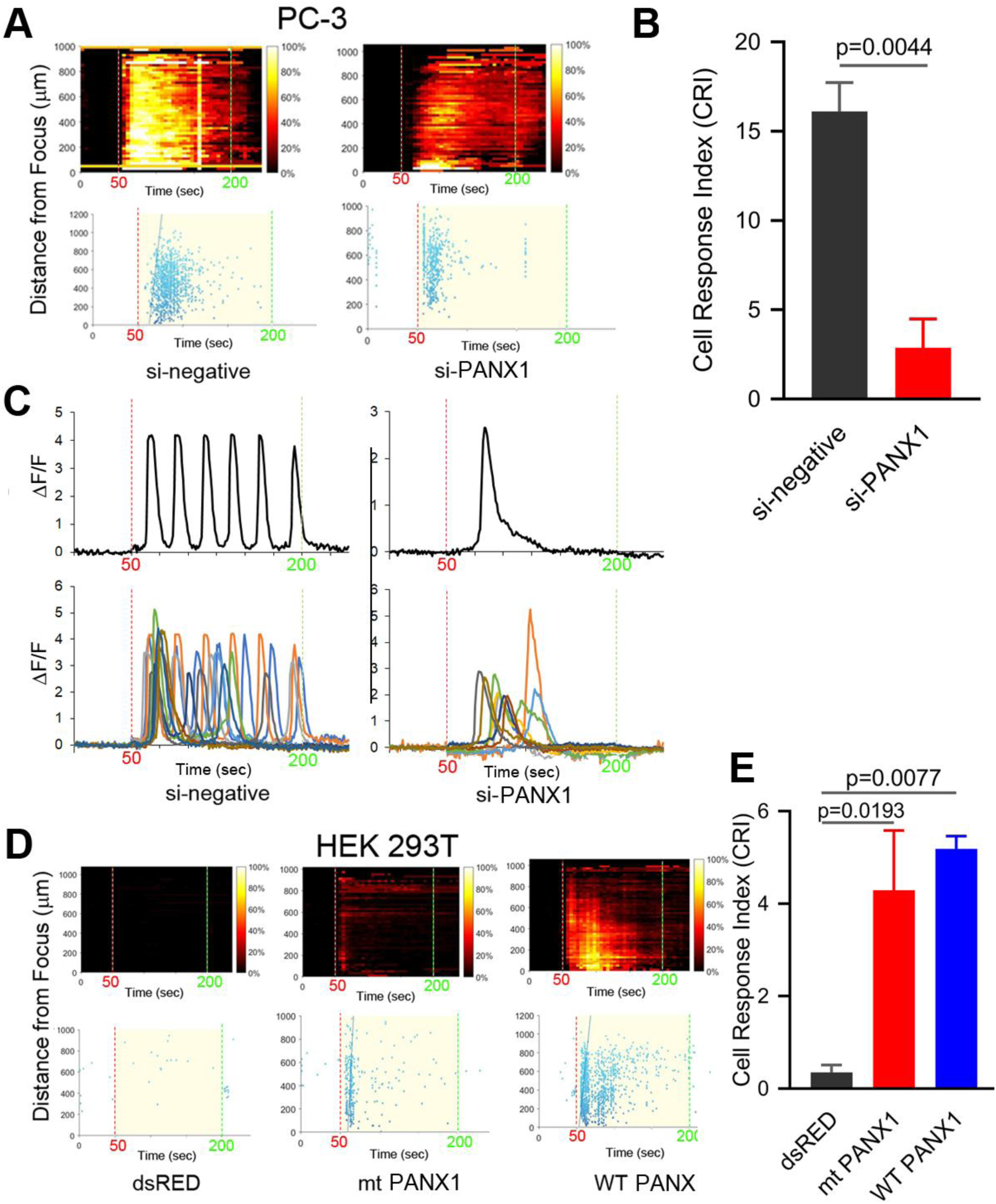
PANX1 expression appears to be both necessary and sufficient for intracellular Ca^2+^ responses. **(A)** si-PANX1 RNA treatment in PC3 cells reduced Ca^2+^ responses compared to si-negative RNA (scramble) as a control. **(B)** Quantitative cell response index (CRI) values of the si-PANX1 RNA treatments relative to the control. *n*=3. Error bars, s.e.m., exact p values by a two-tailed t-test. **(C)** Fluorescence patterns in cells that first responded to the FUS stimulus after the treatments. One representative cell (top) is shown with fluorescence patterns in ten cells (bottom). **(D)** HEK293T cells transfected with WT PANX1 or mt PANX1^1-89^-mRFP (mt PANX1-mRFP) constructs showed Ca^2+^ responses while control HEK cells transfected with dsRED construct have no FUS-induced Ca^2+^ response. **(E)** Quantitative CRI values of the transfected cells. *n*=3. Error bars, s.e.m., ANOVA, Dunnet’s correction, exact p values.

### PANX1 localizes to ER and PM in PC-3 cells

We next established the cellular localization of PANX1 in PC-3 and HEK cells used in this study. HEK cells transfected with either a WT PANX1-EGFP fusion construct (WT PANX1-EGFP; (Furlow et al., 2015)) or mt PANX1-mRFP construct (**Figure 5A**) and imaged with wide field fluorescence microscopy. The mt PANX1-mRFP fluorescence is detected preferentially in perinuclear regions, consistent with expected ER localization (Furlow et al., 2015) while WT PANX1-EGFP fluorescence localized primarily to PM with a reduced signal localized to putative ER (**Figure 5C**). A similar result was obtained after staining PC-3 cells with anti-PANX1 antibodies that specifically recognize the N and C terminal located epitopes (**Figure 5C**). These observations, together with other studies (Furlow et al., 2015; Vanden Abeele et al., 2006), suggest that the C-terminus of PANX1 is important for PM localization and its absence results in mt PANX1-mRFP accumulating in ER. To further confirm this differential localization, we used high resolution total internal reflectance fluorescence (TIRF) microscopy of transfected HEK cells. WT PANX1-EGFP fluorescence is clearly detected by TIRF at PM while cells expressing mt PANX1-mRFP display little or no PM fluorescence (**Figure 5E**, the first column). Imaging the same cells using wide-field microscopy, both WT PANX1-EGFP and mt PANX1-mRFP signals are observed in the ER (**Figure 5E**, the middle columns). We conclude that WT PANX1-EGFP localizes to both ER and the PM, while mt PANX1-mRFP localization is restricted to ER. Combining this localization data with FUS-dependent stimulus data suggest that FUS may be capable of directly or indirectly stimulating ER-localized PANX1 to evoke the internal Ca^2+^ oscillations.

**FIGURE 5.**
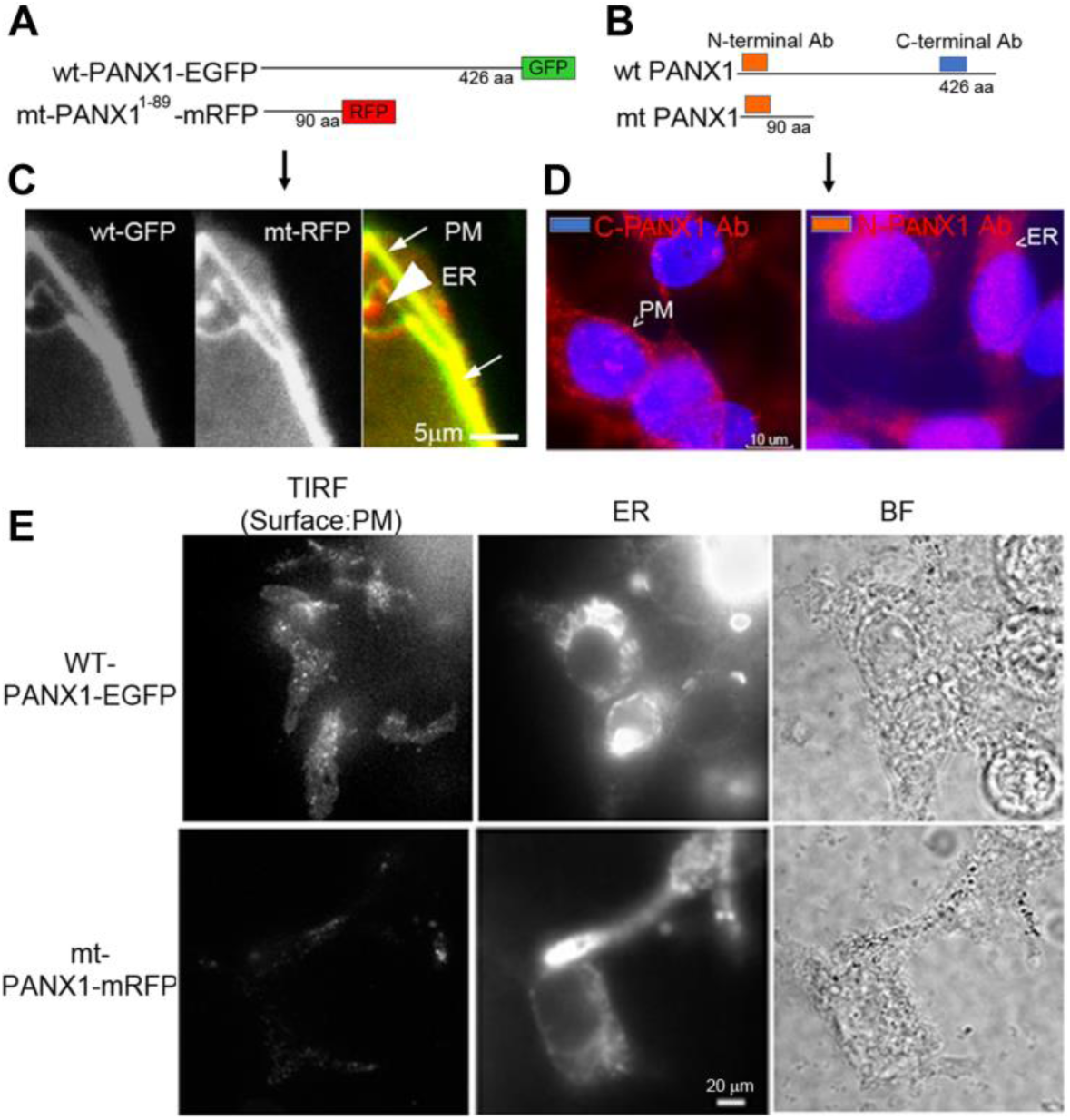
Localization of PANX1. **(A, B)** Schematic of fluorescent WT and mt PANX1 constructs (**A**) and the N and C-terminal epitopes recognized by anti-PANX1 antibodies (Ab) (**B**). **(C)** Localization of WT PANX1-EGFP and mt PANX1-mRFP in transfected HEK cells. **(D)** Localization of endogenous PANX1 in PC-3 cells using N- or C-terminal specific Abs. Nuclear DAPI stain is depicted as blue. **(E)** TIRF imaging on HEK cells transfected by WT PANX1-EGFP or mt PANX1-mRFP constructs. WT PANX1-EGFP localizes in the PM and the ER, while mt PANX1-mRFP only in the ER.

### FUS-dependent Ca^2+^ oscillatory response does not depend on cytoskeletal integrity

The cytoskeleton is believed to be important for the transmission of mechanical forces to internal cellular structures (Cox et al., 2016; Fletcher and Mullins, 2010) following US stimulation of mechanosensitive PM channels. P ANX1 channels are mechanosensitive (Bao et al., 2004). We tested the role of cytoskeletal integrity in FUS stimulation by addition of cytoskeletal protein/process disrupters, including CytochalasinD (actin filaments), Nocodazole (microtubules), ML-7 and Blebbistatin (actomyosin contractility). FUS-evoked Ca^2+^ oscillatory responses appeared essentially normal in PC-3 cells when any of these disruptors were present (**Figure 6A-C**) and are thus not dependent on intact functional cytoskeletal proteins in invasive PC-3 cells. Additionally, this result suggests that FUS may be able to directly mechanostimulate ER localized PANX1. This result is in contrast with previous studies that identified an important role for cytoskeletal networks in transducing US stimuli (De Cock et al., 2015). An important difference between our high frequency non-contact focused US stimulus and most other studies is that the latter uses low frequency US and requires physical contact with the PM through microbubbles (Burks et al., 2019; Carreras-Sureda et al., 2018; Clapham, 2007). This could explain why an intact cytoskeleton appeared necessary for internal Ca^2+^ release in these other studies. We conclude from our results that FUS stimulation appears to be sufficient to result in internal mechanosensory activation of ER localized PANX1 and this coupling results in Ca^2+^ release from internal stores.

**FIGURE 6.**
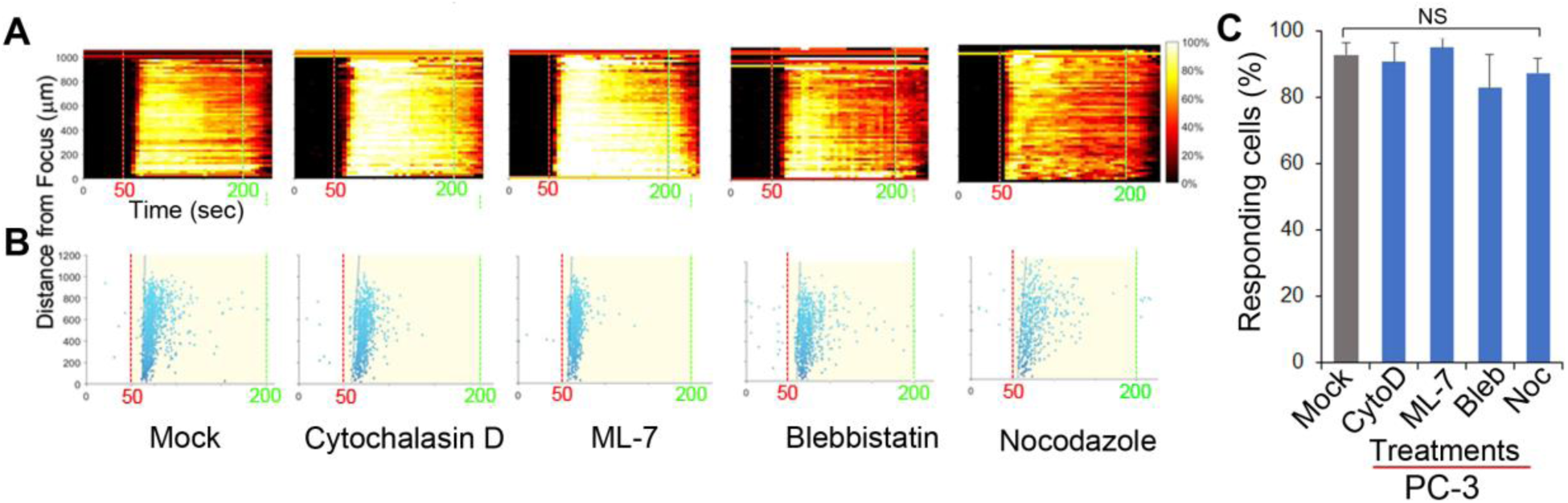
Effect of inhibitors of cytoskeletal support and actomyosin on FUS-induced Ca^2+^ responses in PC-3 cells. The responses are represented when cells were treated with mock, 2 μM CytochalasinD (CytoD), 5 μM ML-7, 5 μM Blebbistatin (Bleb), and 1 μM Nocodazole (Noc). **(A)** The percentage of responding cells over time. **(B)** The time at which each cell first responded to the stimulus. **(C)** Percentage of responding cells after the treatments. Cytoskeletal support and actomyosin did not affect FUS-induced calcium responses. None of them reduced the calcium responses, suggesting a distinctiveness of FUS. *n*=4. Error bars, s.e.m., NS, not significant, by a one-tailed t-test.

### FUS stimulation induces propagation of intercellular Ca^2+^ waves mediated via ATP release and PANX1

We previously demonstrated that FUS-induced calcium waves were not caused by ultrasonic surface waves or gap junction-mediated paracrine signaling (Weitz et al., 2017). However, they may depend on paracrine as well as autocrine signaling via the release of extra-cellular messengers, such as ATP. To test this possibility, we performed FUS stimulation of PC-3 cells in the presence of extracellular apyrase (an ATP degrading enzyme) or in the presence of Suramin or PPADS (2 purinergic receptor blockers). These treatments completely abolished FUS-stimulated Ca^2+^ responses (**Figure 7A-D**). These data indicate that extracellular ATP can induce Ca^2+^ waves, and that FUS stimulation might evoke ATP release into the extracellular space where it activates PM-bound purinergic receptors on the same or nearby cells (e.g. P2X or P2Y). It is unlikely, however that P2×7 or P2Y1 receptors are involved in this process due to pharmacological studies summarized in **Table1** and **Figure S3**.

**FIGURE 7.**
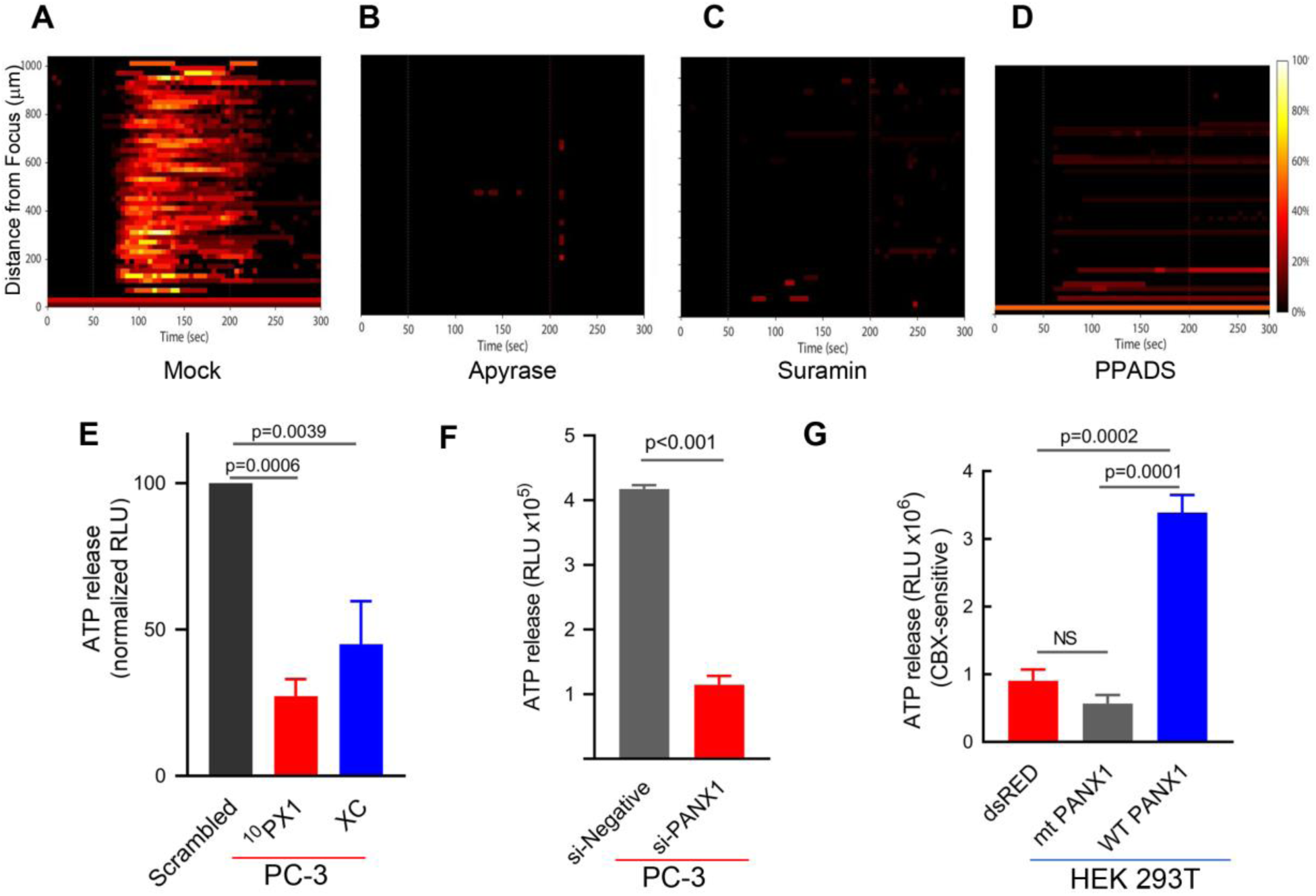
Effects of intercellular Ca^2+^ wave inhibitors on Ca^2+^ response (A-D), and effects of PANX1 modulation on ATP release (E-G). The responses are represented when PC-3 cells were treated with Apyrase, Suramin, PPADS and mock; Mock **(A)**, 20-50 units/ml Apyrase **(B)**, 100 μM Suramin **(C)**, and 100 uM PPADS **(D)**. The percentage of responding cells over time was shown. Experimental results presented are representative and were independently replicates at least two times with three independent biological samples. **(E)** Quantification of PANX1-mediated ATP release from PC-3 cells pretreated for 15 min with scrambled (SC), ^10^PX1, or Xestospongin C (XC). *n*=4. ANOVA, Dunnet’s correction, exact p values. **(F)** Quantification of PANX1-mediated ATP release from PC-3 cells transfected with control si-negative or si-PANX1 RNA. *n*=4, p values by a two-tailed t-test. **(G)** Quantification of PANX1-mediated ATP release from HEK cells transfected with dsRED, mt PANX1-mRFP or WT PANX1-EGFP, and pretreated for 10 min with CBX (500uM). *n*=3. ANOVA, Dunnet’s correction, exact p values. Error bars, s.e.m., NS, not significant.

In our previous study we showed that FUS stimulation and subsequent Ca^2+^ responses did not depend on formation of gap junctions (Weitz et al., 2017). This finding supports that PANX1 forms ATP permeant hemichannels which mediate extracellular Ca^2+^ wave propagation. To evaluate this, we treated PC-3 cells with ^10^PX1 or si-*PANX1* and measured ATP levels. Treated cells have significantly reduced extracellular ATP release (**Figure 7E, F**), indicating that PC-3 cells mediate substantial ATP release through PANX1 channels.

To determine whether mt PANX1-mRFP or WT PANX1-EGFP alters extracellular ATP release through PANX1 channels, we measured CBX-sensitive extracellular ATP release from HEK cells expressing mt PANX1-mRFP or WT PANX1-EGFP. PANX1-mediated ATP release was quantified by measuring the reduction in ATP release in the presence of CBX (Chekeni et al., 2010; Gulbransen et al., 2012; Thompson et al., 2008). When WT PANX1-EGFP was expressed in HEK cells, CBX-sensitive ATP release was enhanced (**Figure 7G**). However, CBX-sensitive ATP release was not enhanced when mt PANX1-mRFP was expressed (**Figure 7G**). This suggests that mt PANX1-mRFP, localized to ER is capable of mediating intracellular Ca^2+^ release (see **Figure 4D, E**) but that it may operate differently than PM WT PANX1. Perhaps mt PANX1 lacking the C-terminal amino acids cannot form homo-oligomers which are necessary to form functional PM ATP release channels (Romanov et al., 2012; Wang et al., 2014; Wang and Dahl, 2018). Additional work will be necessary since the construct we used also contains and additional RFP fusion which may interfere with oligomer formation.

### FUS stimulation induces cytokine/chemokine secretion from PC-3 cells

PANX1 is important for inflammasome activation (Silverman et al., 2009). Using a human cytokine array, we examined whether FUS effectively triggers PC-3 cells to secrete specific cytokines and chemokines. The assay was performed on the supernatants of cells grown for 1 day in media supplemented with charcoal-stripped fetal bovine serum after 46-MHz FUS stimulation repeated 5 times under several conditions (**Figure 8A**). PC-3 cells exhibited Ca^2+^ responses over a range of ultrasound intensities (∼300-1,155 mW/cm^2^) (**Figure 1E-G**), while non-invasive BPH-1 cells showed no response at this range of intensities (Weitz, 2017). Notably, FUS stimulation showed both qualitative and quantitative differences in the levels of cytokine and chemokine secretion from PC-3 cells as the intensity of stimulation was varied (**Figure 8B**). This suggests that FUS stimulation may be fine-tuned to control release of specific cytokine/ chemokine profiles, an exciting possibility with potentially important therapeutic applications.

**FIGURE 8.**
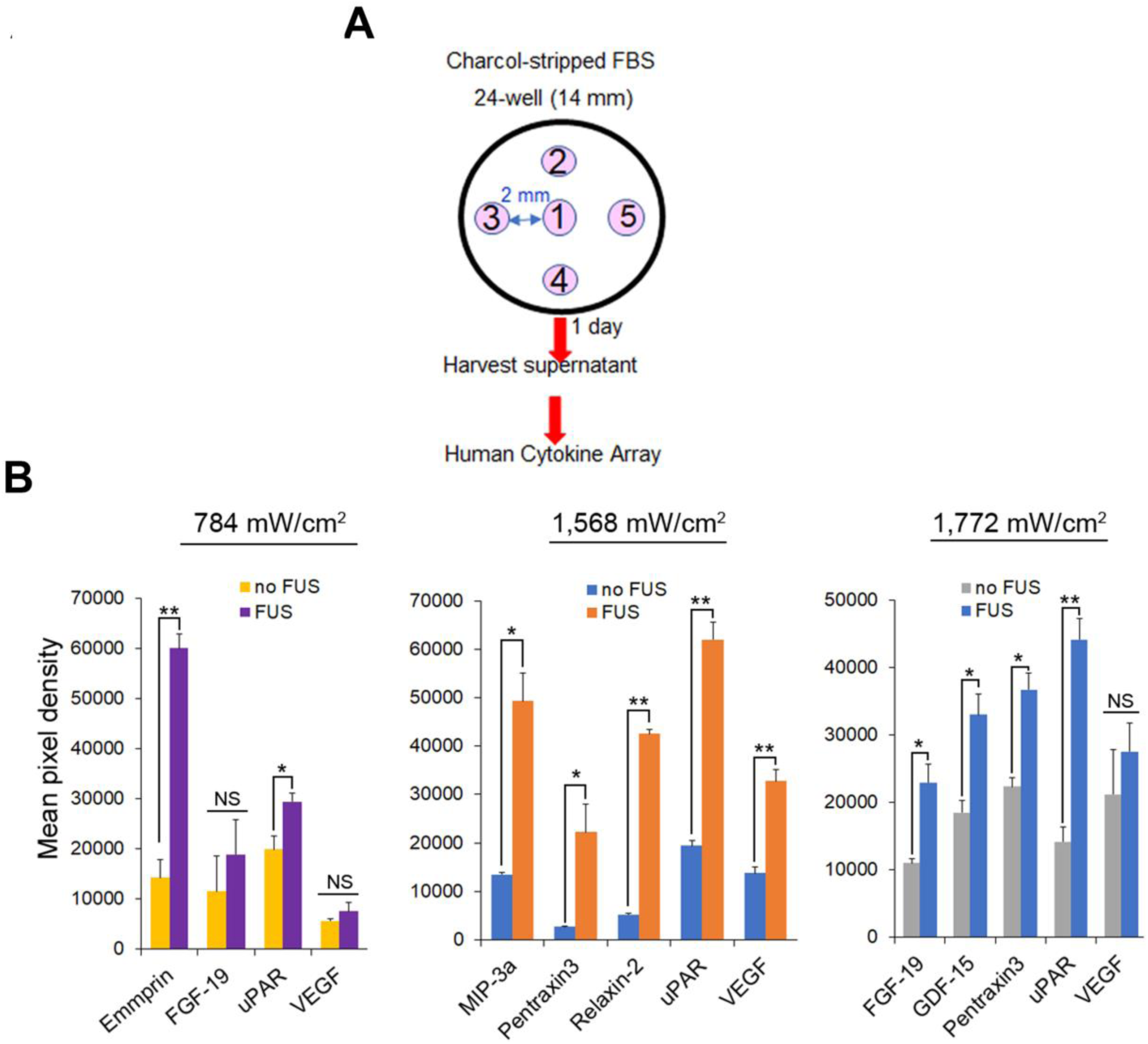
Cytokine and chemokine secretion from PC-3 cells following different intensity of FUS stimulation. **(A)** Protocol for FUS stimulation and a human cytokine array. **(B)** Cytokine and chemokine secretion from PC-3 cells following different intensity of FUS stimulation. *n*=2. Error bars, s.e.m., NS, not significant, *, P<0.05; **, P<0.01 by a one-tailed t-test.

## DISCUSSION

Our previous work demonstrated that mechanosensory FUS-stimulation generates a robust Ca^2+^ signaling response which can be used to distinguish invasive from non-invasive cancer cells (Hwang et al., 2013; Weitz et al., 2017). Others have also demonstrated mechanosensory Ca^2+^ signaling responses using non-focused US-stimulation other contexts (Carina et al., 2018; Pan et al., 2018; Wood and Sehgal, 2015) as well as more general studies to clarify the physiological, cell biological and molecular mechanisms underlying mechanosensory dependent Ca^2+^ signaling responses (Castellanos et al., 2016; Liu et al., 2017; Maresca et al., 2018; Tyler et al., 2008). Our results include several new findings that further clarify the mechanically responsive Ca^2+^ signaling pathways and identify a new role for PANX1 in mediating the FUS-dependent responses.

Removing or lowering external Ca^2+^ from culture medium did not eliminate FUS dependent Ca^2+^ signaling (see **Figure 1D**). This raised the possibility that an internal mechanosensory event is present and coupled to Ca^2+^ release from an internal storage site. Other work has also identified an ER dependent Ca^2+^ response mechanism using more conventional US stimulation involving IP_3_R activation (Burks et al., 2019). This response required an intact cytoskeleton believed to be important for the mechanotransduction of the stimulus to the ER membrane localized IP_3_R (Kim et al., 2015). However, in our study, FUS-dependent internal Ca^2+^ release is present even when cytoskeletal integrity has been disrupted (see **Figure 6**). This raises the interesting possibility that FUS mechanostimulation may be able to directly activate internal Ca^2+^ release and we identified mechanosensitive PANX1, partially localized to ER, as the potential internal target for this process (see **Figures 3-5**).

PANX1 is localized to the PM where it functions as an ATP release channel involved in intercellular signaling events (Wang and Dahl, 2018). We also confirm this for PC-3 cells (see **Figure 7E-G**). PANX1 channels respond to different types of chemical and mechanical stimuli with distinct channel open conformations (‘large’ and ‘small’) (Dahl, 2018). CBX and PB inhibit both PANX1 conformations (Dahl, 2018). In our study Ca^2+^ responses are eliminated in PC-3 cells treated with CBX or PB (**Table 1, Figure S4**) suggesting that these cells contain both PANX1 conformers.

The “small” conformer of PANX1 channel is reported to be impermeant to ATP (Romanov et al., 2012; Wang et al., 2014; Wang and Dahl, 2018). We show that HEK cells transfected with mt PANX1-mRFP also do not release ATP but can confer the internal Ca^2+^ response (see **Figure 4**). Perhaps the mt PANX1-mRFP construct we used is functionally similar to the “small” form of PANX1. Interestingly, a mutant truncated PANX1 channel (PANX1^1-89^) has also been associated with highly metastatic breast cancer cells (Furlow et al., 2015). Taken together these results suggest that FUS-induced ER calcium release mediated through mt PANX1 may play a key role in cancer cell invasion and tumor metastasis. Further studies will be required to determine the mechanistic significance of this correlation.

While PANX1 has clearly been localized to ER (Furlow et al., 2015; Vanden Abeele et al., 2006), see also **Figure 5**) its functional significance is largely unknown. In addition to demonstrating its potential role in Ca^2+^ release from ER, we show a remarkably similarity with some aspects of previously established IP_3_R function in this regard (Mery et al., 2005; Rizaner et al., 2016; Sasse et al., 2007) (see **Figure 3**). Since the simultaneous inhibition of IP_3_R and PANX1 exhibit similar patterns of FUS-dependent internal Ca^2+^ release they may be part of the same Ca^2+^ signaling pathway and provide a mechanism to transduce various stimuli into similar cellular responses.

At high intensities, FUS has been used clinically to thermally ablate tumor cells (FUS and Cancer Immunotherapy, Workshop, 2019). Perhaps more importantly at lower intensities FUS has been shown to stimulate an inflammatory response cancer models which can boost the efficacy of immunotherapy (Bonaventura et al., 2019; Curley et al., 2017; Li et al., 2018; Mauri et al., 2018). Low-intensity FUS, has not yet used in cancer therapy, partly due to our limited understanding of its effects and mechanism of action. In our study we clearly show the potential to use “tuned” FUS to specifically control release of different cytokine/chemokine profiles from invasive cancer cells since this varies as the amplitude of the stimulation was changed (see **Figure 8C**). We are now attempting to extend this exciting observation to better understand FUS-induced anti-tumor immune response modulation by linking it to specific signaling pathways/ molecules and epigenetic dynamics in simple cellular cancer models. This proof of principle work is critical before proceeding to in vivo experimentation or clinical utility.

For example, FUS applied to tumors could potentially modulate immune responses such as the ability to enhance infiltration of tumor targeting CAR T cells. Immunologically “cold” tumors are cancers that contain few infiltrating T cells thus making them impervious to current immunotherapy treatments (Bonaventura et al., 2019; Li et al., 2018). Classically immunologically “cold” cancers include glioblastomas, ovarian, prostate, pancreatic, and most breast cancers, all extremely resistant to current therapies. FUS can be potentially be used as an adjunct therapy to induce secretion of cytokines/chemokines from ‘cold’ cancer cells and mediate conversion into a ‘hot’ tumor responsive to immunotherapy (Curley et al., 2017; Mauri et al., 2018). Of course, more work will be required in a well-controlled cellular model to understand the critical signaling pathways/ molecules and mechanisms necessary for successful clinical translation of this technology.

FUS produces a focused beam of acoustic energy that precisely and accurately reaches large targets in the body without damaging surrounding normal cells (Mittelstein et al., 2020). One of the most striking findings of our study is the suggestion that FUS may also directly stimulate intracellular mechanosensory proteins located on particular membrane limited organelles such as ER. This rases the possibility that future studies could be designed to prove this by designing appropriate reporter constructs, i.e. sensors and bioswitches (Kim et al., 2015; Piraner et al., 2017), and optimizing stimulus parameters (e.g. amplitude, frequency, duty factor and duration) and thus provide a new tool to study mechanosensitive intracellular processes.

In summary, we demonstrate that non-contact mediated FUS stimulates ER localized PANX1 to initiate an intracellular Ca^2+^ release. This process does not require an intact cytoskeleton and is independent of external Ca^2+^entry. PM localized PANX1, however, does appear necessary to mediate the intercellular spreading of Ca^2+^ waves likely through ATP release. In addition, FUS stimulation results in the release of specific chemokine/cytokine profiles from invasive PC-3 cancer cells. The specific cytokine/chemokine profile can be modified by varying FUS stimulus intensity.

Taken together our results suggest a new mechanistic working model for FUS-stimulation dependent Ca^2+^ signaling in cells which is shown schematically in **Figure 9A**. The initial cytoplasmic Ca^2+^ signal subsequently results in extracellular ATP release, possibly mediated by PM-PANX1 action and/or direct FUS stimulation. The ATP acts on purinergic receptors in nearby cells, thus propagating the spread of intercellular Ca^2+^ waves. Overall, these processes are not dependent on cytoskeletal integrity or other types of Ca^2+^ channels present in ER. The initial ER Ca^2+^ release, however, is not strictly related to mechanosensory stimulation of ER localized PANX1 but may also be influenced by ER localized IP_3_Rs as reported by others (Bootman et al., 2002; Diver et al., 2001; Xu et al., 2005). An additional response of FUS stimulation of PC-3 invasive cancer cells is the coupled release of specific cytokines/chemokines release from PC-3 cells. This new model can be compared to current working model largely derived from conventional US stimulation for comparison (**Figure 9B**) (Burks et al., 2019; Carreras-Sureda et al., 2018; Clapham, 2007).

**FIGURE 9.**
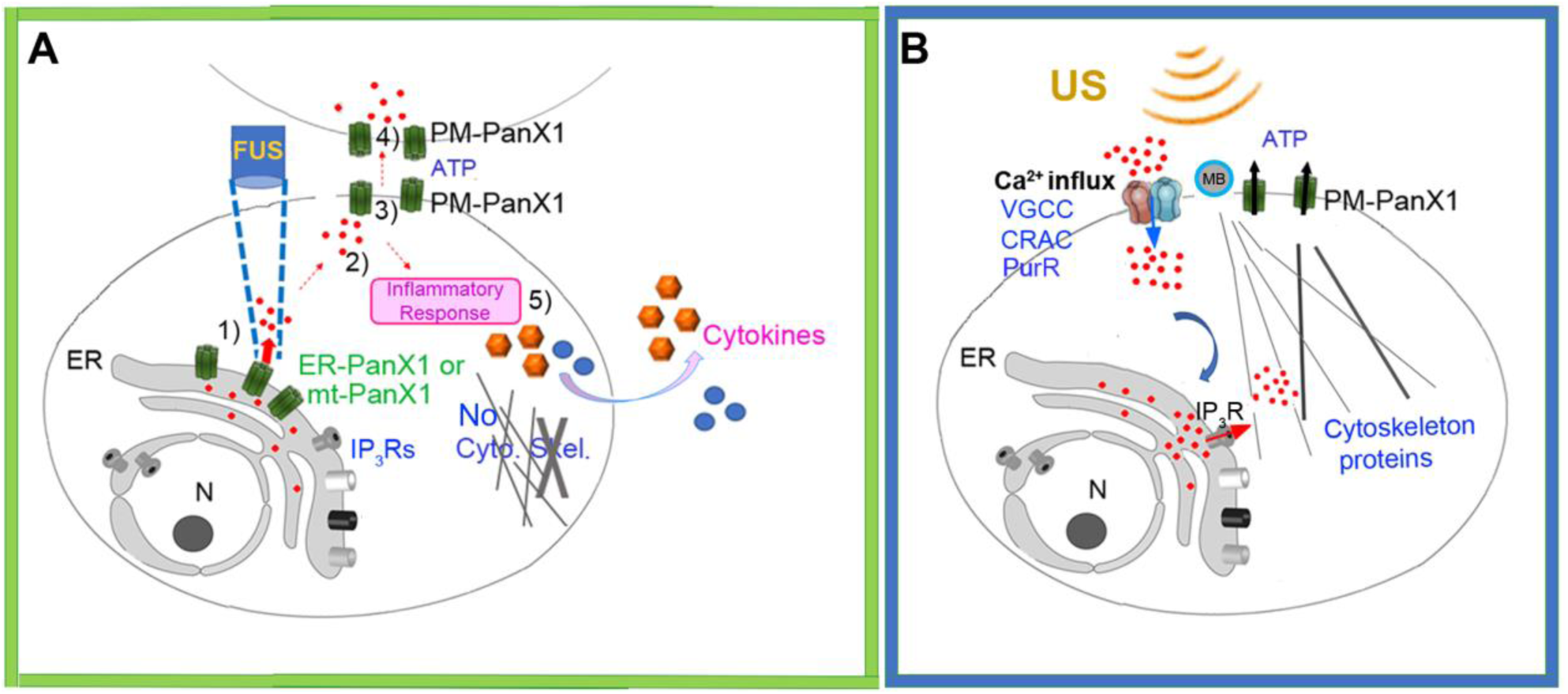
(A). Schematic of new working model FUS-dependent response mechanisms in PC-3 invasive cancer cells. 1) FUS stimulation activates ER localized mechanosensitive PANX1 resulting in internal Ca^2+^ release from ER stores. 2) This cytoplasmic Ca^2+^ signal stimulates ATP release through PANX1 PM channels. 3) The released ATP acts on purineregic receptors, many in adjacent cells. 4) This results in a propagating extracellular Ca^2+^ wave which spreads through the cell population possibly via PM PANX1 or opening of PM Ca^2+^ channels. 5) FUS stimulation also results in secretion of cytokines/chemokines. **(B)** Schematic of currently accepted working model based largely on conventional US stimulation with no proposed role for internal ER Ca^2+^ release but rather a link to US energy transduction to ER mediated by IP_3_R. MB, Microbubbles.

## AUTHOR CONTRIBUTIONS

N.S.L. conceived and designed experiments. N.S.L. and P.M.S. wrote the manuscript. C.W.Y., Q.W., S.M. and K.M.K. executed Ca^2+^ imaging. C.W.Y. and Q.W. analyzed the Ca^2+^ imaging. N.S.L. executed all molecular and cell biological experiments and assays with C.W.Y. and Q.W. H.J. (46-MHz) and G.L. (3-MHz) characterized transducers with a needle hydrophone. R.C. and L.J. fabricated transducers in house. A.F. and F.P. performed TIRF imaging. A.C.W. helped develop calcium imaging program. P.M.S., A.C.W., R.H.C and K.K.S contributed helpful discussions/advice. C.W.Y, P.M.S. A.C.W, F.P. and K.K.S. edited the manuscript with helpful suggestions. All authors reviewed the manuscript.

## ACKNOWLEDGEMENTS

We thank Sohail F. Tavazoie (Rockefeller University) for providing WT PANX1-EGFP and mt PANX1-mRFP cDNA constructs.

