## Supplementary material for "Focused ultrasound stimulates ER localized mechanosensitive PANNEXIN-1 to mediate intracellular calcium release in invasive cancer cells": SUPPLEMENTARY MATERIAL.docx

**FIGURE S 1. Effect of FUS stimulation amplitude on PC-3 cell calcium response using 3-MHz transducer.** All stated voltages represent peak-to-peak amplitude (Vp-p). Values in parentheses indicate the mV at each voltage, as measured by a hydrophone. **(A)** 2-D histograms showing the percentage of responding cells over time. **(B)** Scatter plots showing the time at which each cell first responded to the stimulus (each dot represents a responding cell). **(C)** Quantitative percentage of responding cells. *n*=3. Error bars, s.e.m., *n* represents biological replicates.

**FIGURE S 2. Effect of FUS stimulation amplitude on HEK cell calcium response using 46-MHz transducer.** All stated voltages represent peak-to-peak amplitude (Vp-p). Values in parentheses indicate the mV at each voltage, as measured by a hydrophone. Quantitative percentage of responding cells. *n*=3. Error bars, s.e.m., *n* represents biological replicates. Some spontaneous response background was occasionally shown, so the percentages of responding cells are ~10%. However, the calcium response in HEK cells was not altered by different FUS stimulation amplitude, which is different from PC-3 cells.

**FIGURE S 3. Effect of treatment of inhibitors on PC-3 cell calcium response.** 2-D histograms showing the percentage of responding cells over time. (**A)** Effect of treatments of P2 receptor inhibitors on PC-3 cell calcium response**. (B)** Effect of treatments of Ca^2+^ influx inhibitors on PC-3 cell calcium response. These did not change the calcium response.

**FIGURE S 4. Effect of both treatment of ^10^PX1 and XC on PC-3 cell calcium response using 46-MHz transducer. (A)** 2-D histograms showing the percentage of responding cells over time. **(B)** Scatter plots showing the time at which each cell first responded to the stimulus (each dot represents a responding cell). **(C)** Effect of treatments of CBX, PB and FFA. The histograms showed the percentage of responding cells over time. Treatment of CBX, PB or FFA in PC-3 cells abolished Ca^2+^ responses.

**FIGURE S 5.** Quantitative RT-PCR analysis of WT PANX1 transcript expression in PC3 cells transfected two independent siRNAs that specifically target FL PANX1, as described previously; *n*=2. Error bars, s.e.m., si-*PANX1*-N1 (L-018253-00) showed <35% reduction **(A)** so we did not use it. Another si-*PANX1*-N2 (D-018253-02) was used in most experiments and called as ‘si-*PANX1*’ **(B).** The variations of reduction occurred because of cell heterogeneity.

**Table S 1.** Pharmacological agents used to investigate the mechanism of FUS-induced calcium rise in PC-3 cells.

**Video S1.** Calcium responses of strongly invasive PC-3 prostate cancer cells to stimulation with 46-MHz low-intensity FUS. The FUS stimulus onset and offset times are 50 s and 200 s, respectively.

**Video S2.** Calcium responses of non invasive HEK 293T cells to stimulation with 46-MHz low-intensity FUS. The FUS stimulus onset and offset times are 50 s and 200 s, respectively.

**Videos S3-5.** Calcium responses of strongly invasive PC-3 prostate cancer cells after scrambled peptide (SC, **S3**), ^10^PX1 peptide (**S4**) and Xestospongin C (XC, **S5**) application for 20 min, to stimulation with 46-MHz low-intensity FUS. The FUS stimulus onset and offset times are 50 s and 200 s, respectively.

**Videos S6-7.** Calcium responses of strongly invasive PC-3 prostate cancer cells after si-negative (**S6**) or si-*PANX1* (**S7**) treaments for 2 days, with 46-MHz low-intensity FUS. The FUS stimulus onset and offset times are 50 s and 200 s, respectively.

**Videos S8-10.** Calcium responses of non invasive HEK 293T cells, after transfection of dsRED (**S8**), mt PANX1-mRFP (**S9**) or WT PANX1 (**S10**) constructs, to stimulation with 46-MHz low-intensity FUS. The FUS stimulus onset and offset times are 50 s and 200 s, respectively.

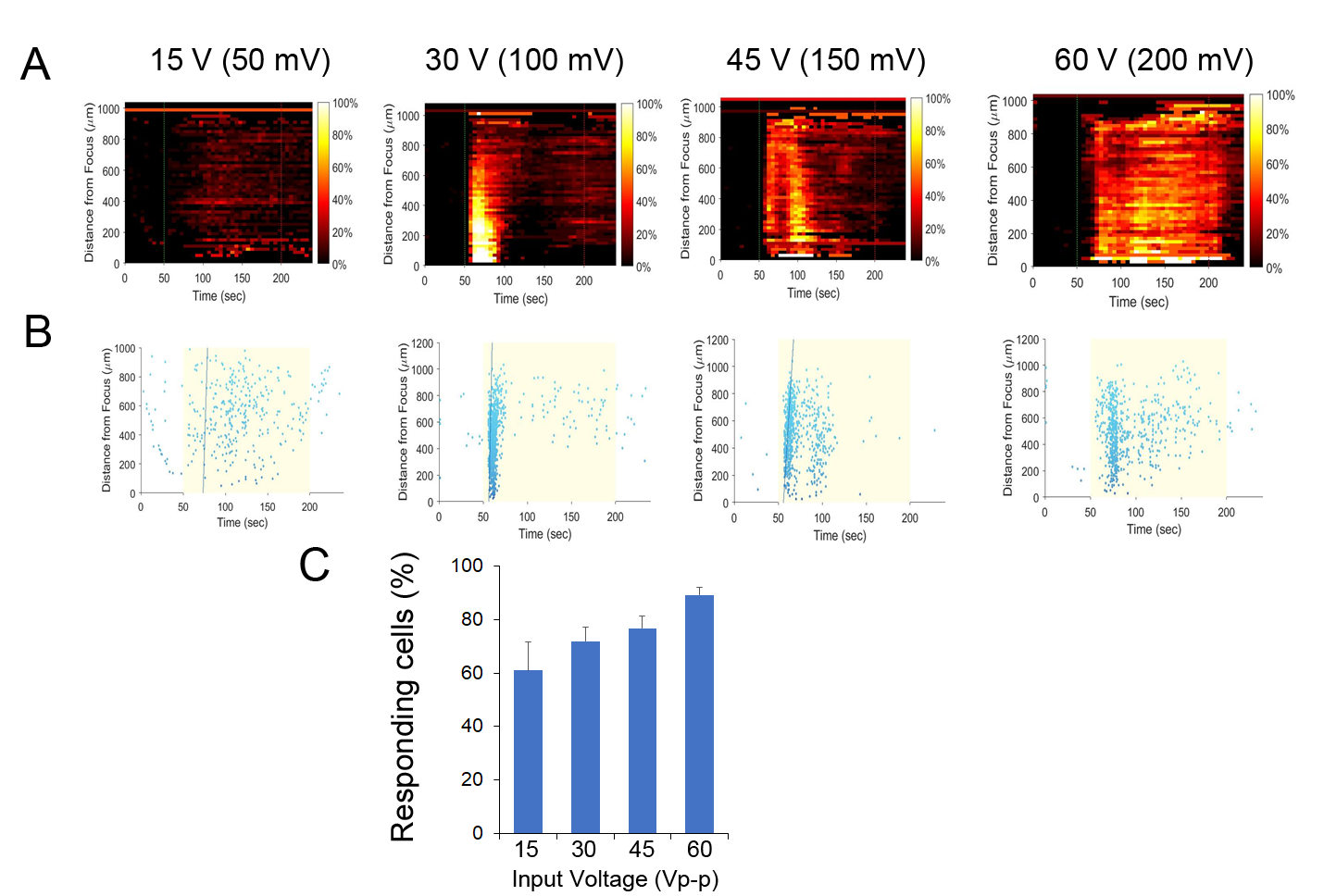

**FIGURE S 1.**  **Effect of FUS stimulation amplitude on PC-3 cell calcium response using 3-MHz transducer.** All stated voltages represent peak-to-peak amplitude (Vp-p). Values in parentheses indicate the mV at each voltage, as measured by a hydrophone. **(A)** 2-D histograms showing the percentage of responding cells over time. **(B)** Scatter plots showing the time at which each cell first responded to the stimulus (each dot represents a responding cell). **(C)** Quantitative percentage of responding cells. *n*=3. Error bars, s.e.m., *n* represents biological replicates.

.

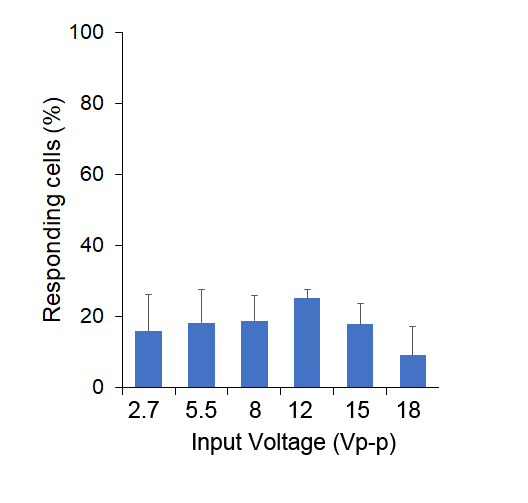

**FIGURE S 2. Effect of FUS stimulation amplitude on HEK cell calcium response using 46-MHz transducer.** All stated voltages represent peak-to-peak amplitude (Vp-p). Values in parentheses indicate the mV at each voltage, as measured by a hydrophone. Quantitative percentage of responding cells. *n*=3. Error bars, s.e.m., *n* represents biological replicates. Some spontaneous response background was occasionally shown, so the percentages of responding cells are ~10%. However, the calcium response in HEK cells was not altered by different FUS stimulation amplitude, which is different from PC-3 cells.

Mock	AZ11645373	MRS2179

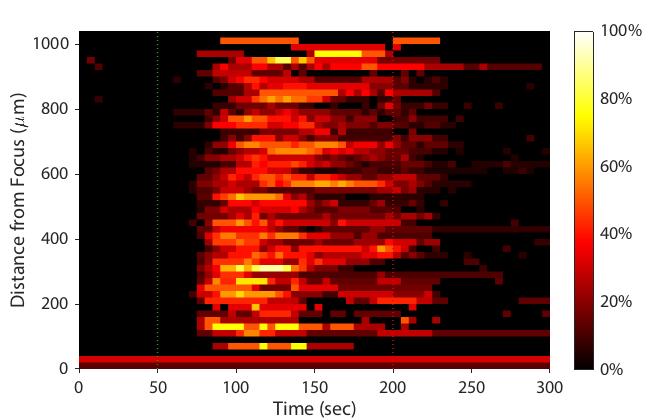

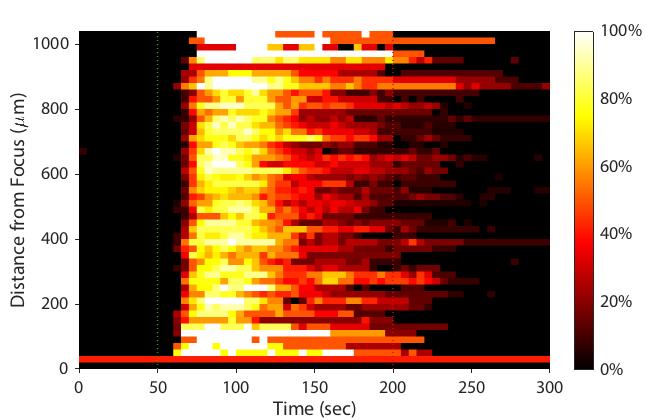

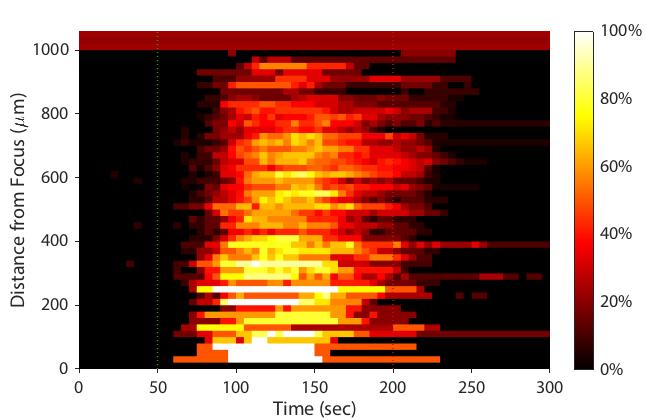

Mock	BTP2	SKF96365

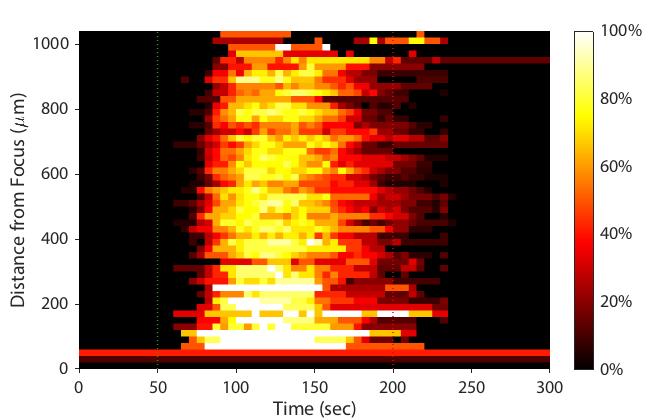

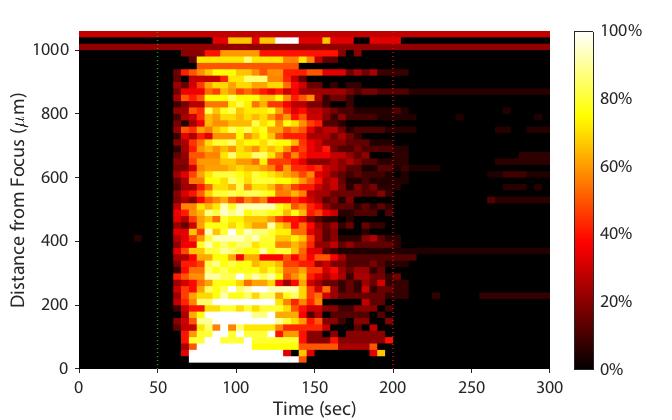

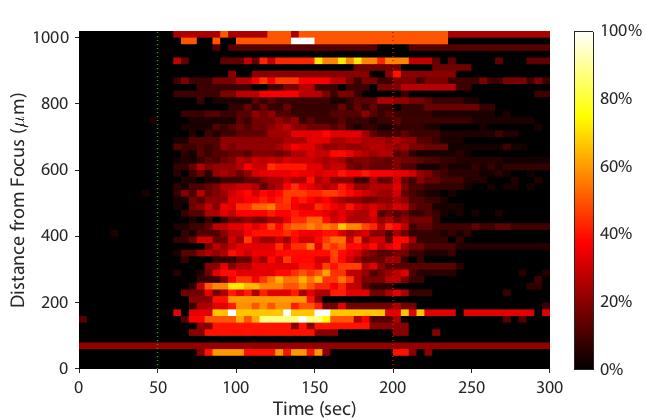

A

B

**FIGURE S 3. Effect of treatment of inhibitors on PC-3 cell calcium response.** 2-D histograms showing the percentage of responding cells over time. (**A)** Effect of treatments of P2 receptor inhibitors on PC-3 cell calcium response**. (B)** Effect of treatments of Ca^2+^ influx inhibitors on PC-3 cell calcium response. These did not change the calcium response.

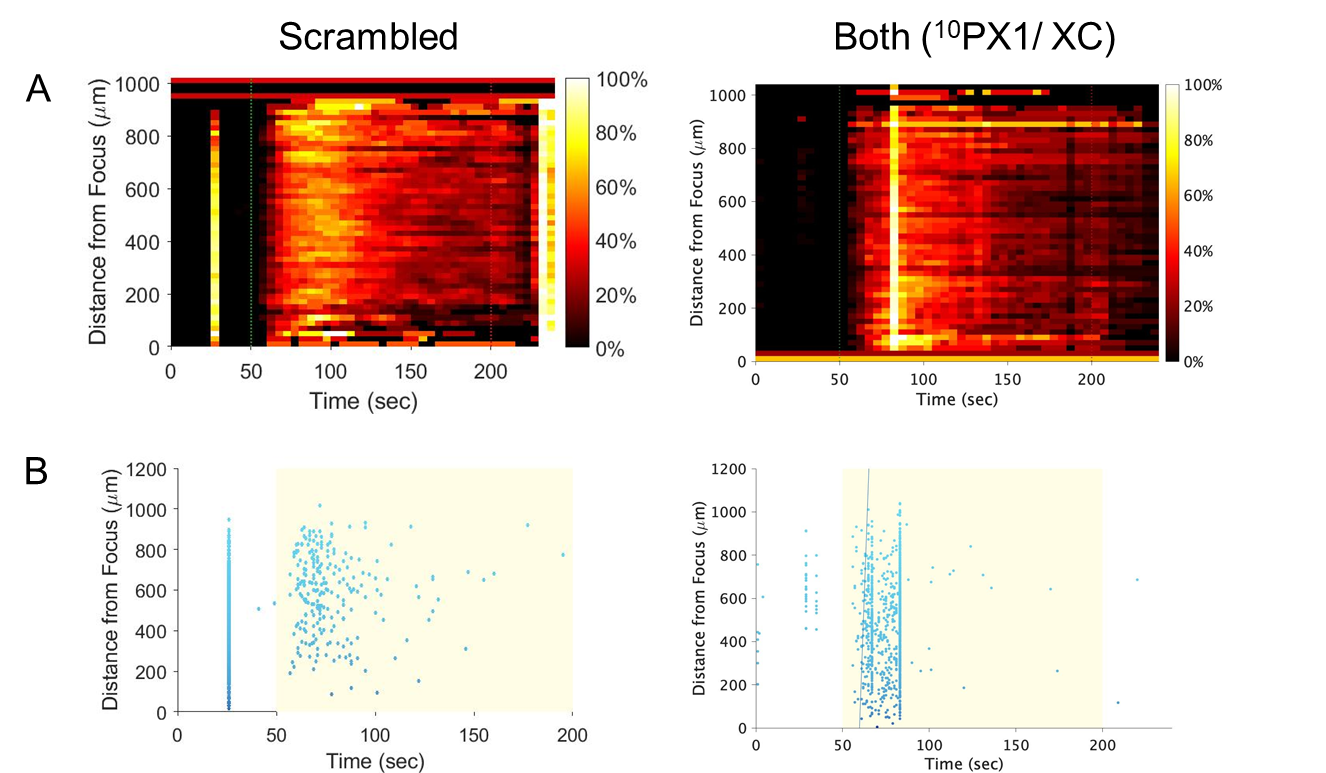

Mock	CBX	PB	FFA

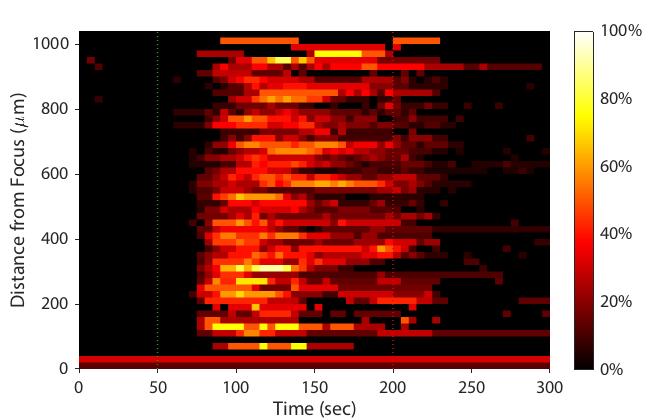

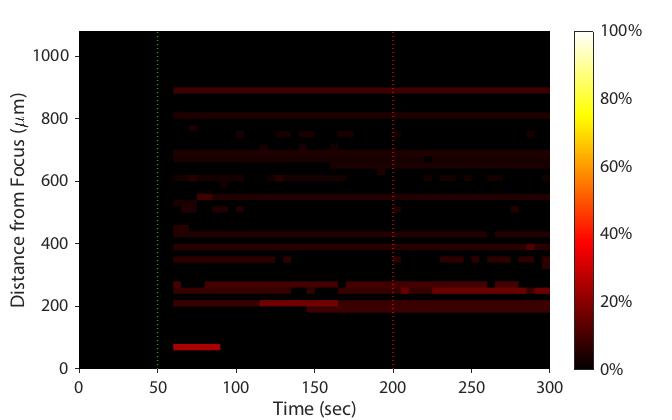

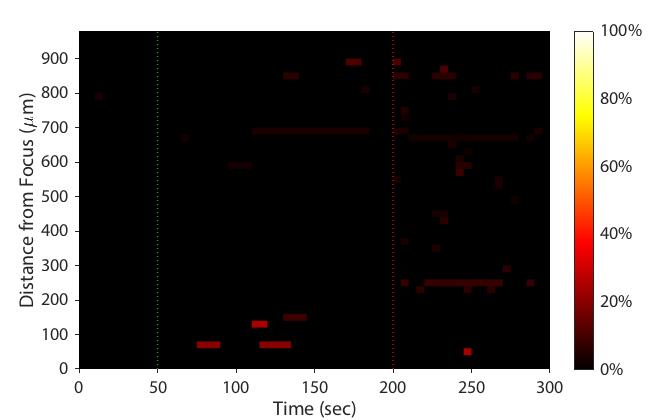

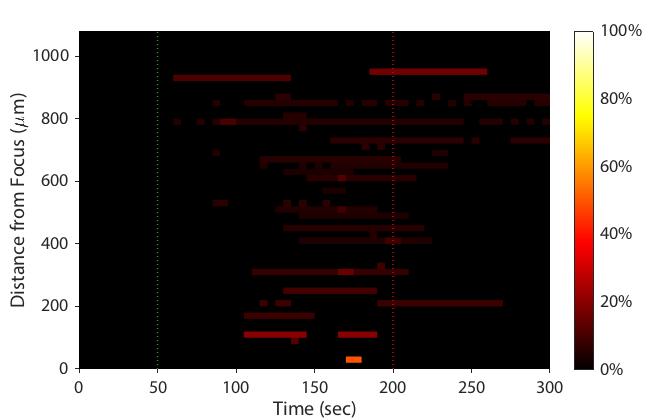

C

**FIGURE S 4. Effect of both treatment of ^10^PX1 and XC on PC-3 cell calcium response using 46-MHz transducer. (A)** 2-D histograms showing the percentage of responding cells over time. **(B)** Scatter plots showing the time at which each cell first responded to the stimulus (each dot represents a responding cell). **(C)** Effect of treatments of CBX, PB and FFA. The histograms showed the percentage of responding cells over time. Treatment of CBX, PB or FFA in PC-3 cells abolished Ca^2+^ responses.

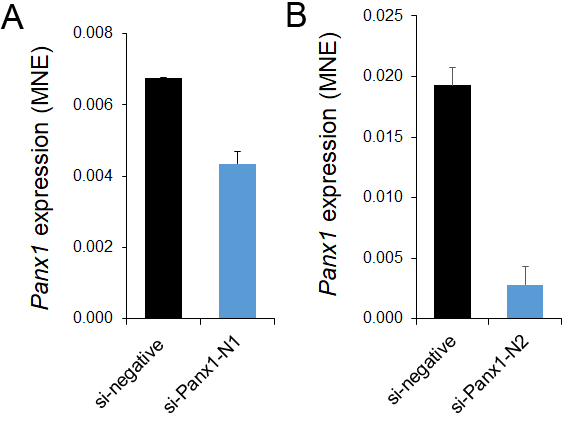

**FIGURE S 5.** Quantitative RT-PCR analysis of WT PANX1 transcript expression in PC3 cells transfected two independent siRNAs that specifically target FL PANX1, as described previously; *n*=2. Error bars, s.e.m., si-*PANX1*-N1 (L-018253-00) showed <35% reduction **(A)** so we did not use it. Another si-*PANX1*-N2 (D-018253-02) was used in most experiments and called as ‘si-*PANX1*’ **(B).** The variations of reduction occurred because of cell heterogeneity.

**Table S 1.** Pharmacological agents used to investigate the mechanism of FUS-induced calcium rise in PC-3 cells.

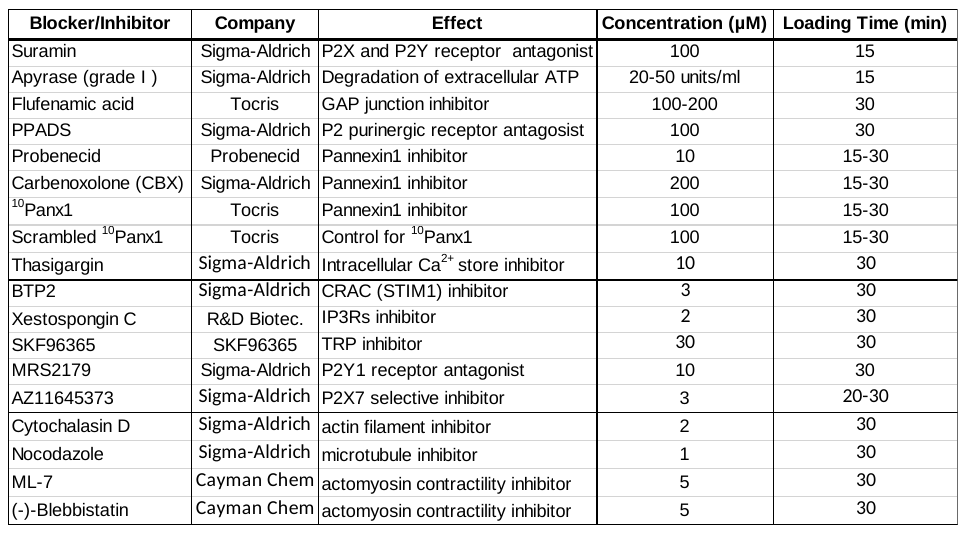
